# Autism sensory dysfunction in an evolutionarily conserved system

**DOI:** 10.1101/297051

**Authors:** Greta Vilidaite, Anthony M. Norcia, Ryan J. H. West, Christopher J. H. Elliott, Francesca Pei, Alex R. Wade, Daniel H. Baker

**Affiliations:** Department of Psychology, Stanford University, CA 94305, USA; Department of Biology, University of York, York, YO10 5DD, UK; Department of Psychology, University of York, YO10 5DD, UK; Department of Psychiatry, Stanford University, CA 94305, USA

**Keywords:** Autism, animal model, *Drosophila*, sensory processing, visual system

## Abstract

There is increasing evidence for a strong genetic basis for autism, with many genetic models being developed in an attempt to replicate autistic symptoms in animals. However, current animal behaviour paradigms rarely match the social and cognitive behaviours exhibited by autistic individuals. Here we instead assay another functional domain – sensory processing – known to be affected in autism to test a novel genetic autism model in *Drosophila melanogaster*. We show similar visual response alterations and a similar development trajectory in *Nhe3* mutant flies (total N=72) and in autistic human participants (total N=154). We report a dissociation between first- and second-order electrophysiological visual responses to steady-state stimulation in adult mutant fruit flies that is strikingly similar to the response pattern in human adults with ASD as well as that of a large sample of neurotypical individuals with high numbers of autistic traits. We explain this as a genetically driven, selective signalling alteration in transient visual dynamics. In contrast to adults, autistic children show a decrease in the first-order response that is matched by the fruit fly model, suggesting that a compensatory change in processing occurs during development. Our results provide the first animal model of autism comprising a differential developmental phenotype in visual processing.

## Introduction

Autism spectrum disorder (ASD) has a strong albeit complex genetic basis with a large number of genes implicated (1–5). A variety of genetic animal models have been proposed for ASD, including murine models (6–8) and more recently, fly models (9). However, for an animal model of any disorder/disease to be useful it needs to fulfill as much face validity as possible (i.e., exhibit a similar phenotype to humans with the disorder/disease). This poses a challenge for multifaceted, heterogenic disorders having symptoms that are difficult to operationalise and measure in animals. While there have been some attempts at measuring defining behaviours of ASD in animal models (10), including difficult to assess social interactions (11), repetitive behaviours (12), and confined interests (13), the links between human symptoms and equivalent animal behaviours are tenuous. For example, social symptoms in mice have been evaluated as defensive behaviour against intruders (11), or as courtship call frequency and wing extension in fruit flies (9), even though neither behaviour manifests in humans.

In addition to the defining social and behavioural features of ASD, autistic individuals report a host of sensory symptoms including unusual sensory interests as well as hyper- and hyposensitivity to intense stimuli such as bright lights or loud noises (14,15). These human ASD sensory processing symptoms have been well documented behaviourally (16–18), with electroencephalography (EEG; 16,17) and neuroimaging (21) and can also be measured in animals using equivalent methods (22). Functioning in sensory systems may be better conserved over evolution than more complex behaviours associated with ASD, therefore we pursued a comparison of sensory responses in humans with ASD and an *Nhe3* fruit fly model of ASD.

A previous study in mice measured visual responses in a related developmental condition, Rett syndrome, and was able to link decreases in visual neural responses and poor visual acuity across species (23). However, it is difficult to generalise these findings to ASD, as human Rett syndrome lacks the pervasive sensory symptoms characteristic of autism (24). An advantageous alternative to rodent models are *Drosophila* given the ease in developing genetic mutations and ability to test many individual animals. Successful *Drosophila* models of human neurological disorders have so far been developed for Parkinson’s disease (25), fragile X syndrome (26) and Alzheimer’s disease (27). Fruit flies share 75% of human disease-causing genes (28) and have a visual system exhibiting similar nonlinear neural properties, including a colour- and luminance-selective module as well as a motion-selective module (29). The neural dynamics of these modules closely resemble those of transient and sustained neural populations in humans (30–32). These factors combine to provide an excellent framework for modelling changes in early sensory neuronal signalling (32) which may lie behind atypical sensory processing in autism.

In this study we evaluated a genetic *Drosophila* model of human ASD by measuring comparable visual responses both in autistic humans and in mutant *Drosophila*. In humans, loss-of-function mutations in the gene *SLC9A9* have been linked to ASD (33). Here we used a *Drosophila* orthologue of *SLC9A9* – *Nhe3*. A homozygous P-element insertion loss-of-function mutants (*Nhe3^KG08307^*) and *Nhe3* hemizygotes *(Nhe3^KG08307^*/Df(2L)BSC187) were used to inhibit *Nhe3* function in fly. The use of two *Nhe3* mutations in different genetic backgrounds ruled out the possibility of other mutations influencing the flies’ visual responses. To assess the functionality of the visual system in these species, we measured steady-state visually evoked potentials (ssVEPs) to temporally-modulated contrast stimuli. During this paradigm a stimulus in flickered on/off at a particular frequency (for example 12Hz) whilst neural responses are recorded from the organism. Using Fourier transformation we then convert time course data into the frequency domain where the amplitude of different frequency components of the neural responses can be measured. From there we extract the 1^st^ harmonic response (which follows the stimulation frequency – 12Hz), as well a 2^nd^ harmonic response. Second harmonics are responses generated by the brightening and darkening transients of the stimulus flicker, thus – 24Hz in the flies and to contrast onset/contrast offset in human. The first and second harmonics probe different aspects of the dynamics of the visual system: sustained and transient neural responses, respectively (34). Previous genetic dissection of the fruit fly has localised the 1^st^ harmonic to photoreceptors and the 2^nd^ harmonic to the lamina (31).

As the visual systems of humans and fruit flies are difficult to compare anatomically, the visual responses obtained here were produced by functionally equivalent human and fruit fly neural substrates. In each organism we assessed the same functional mechanism - contrast transduction. This computation in the fly is performed at the level of photoreceptors and lamina, whereas in humans the same computation is performed in the retina and in early visual cortex (V1). A similar cross-species computational equivalence in the face of vastly different neural substrates has been shown previously for motion perception: third order correlations required for motion perception were found in the lamina of the fly and areas V1 and MT in humans (35).

Furthermore, to investigate the progression of ASD sensory atypicalities over the course of development, we also measured visual responses at two stages of fruit fly maturation and acquired similar responses from autistic children and adults. Finally, as the ASD phenotype is complex and non-binary, we validated our sensory model with a large sample of neurotypical participants with high and low numbers of autistic traits.

## Results

### Increased sustained/transient response ratio in *Nhe3* fruit flies

Using a steady-state visual evoked potential (ssVEP) paradigm (25) (see *Fig 1*) we measured *Drosophila* visual responses to flickering stimuli via an electrode on the fly’s eye. Wild type, eye-colour matched flies (a cross between isogenic and Canton-S) were used as controls (+). Twelve flies from each genotype were tested at three days (when the flies are young; total n=36) and at 14 days post eclosion (older; total n=36). First harmonic (12Hz) and second harmonic (24Hz) response amplitudes were derived by fast Fourier transform (see *Methods*). Although the first harmonic responses of mutant and wild-type flies were the same, the second harmonic response was significantly reduced in the *Nhe3* mutants (*Fig 2a, 2b*).

**Fig 1.**
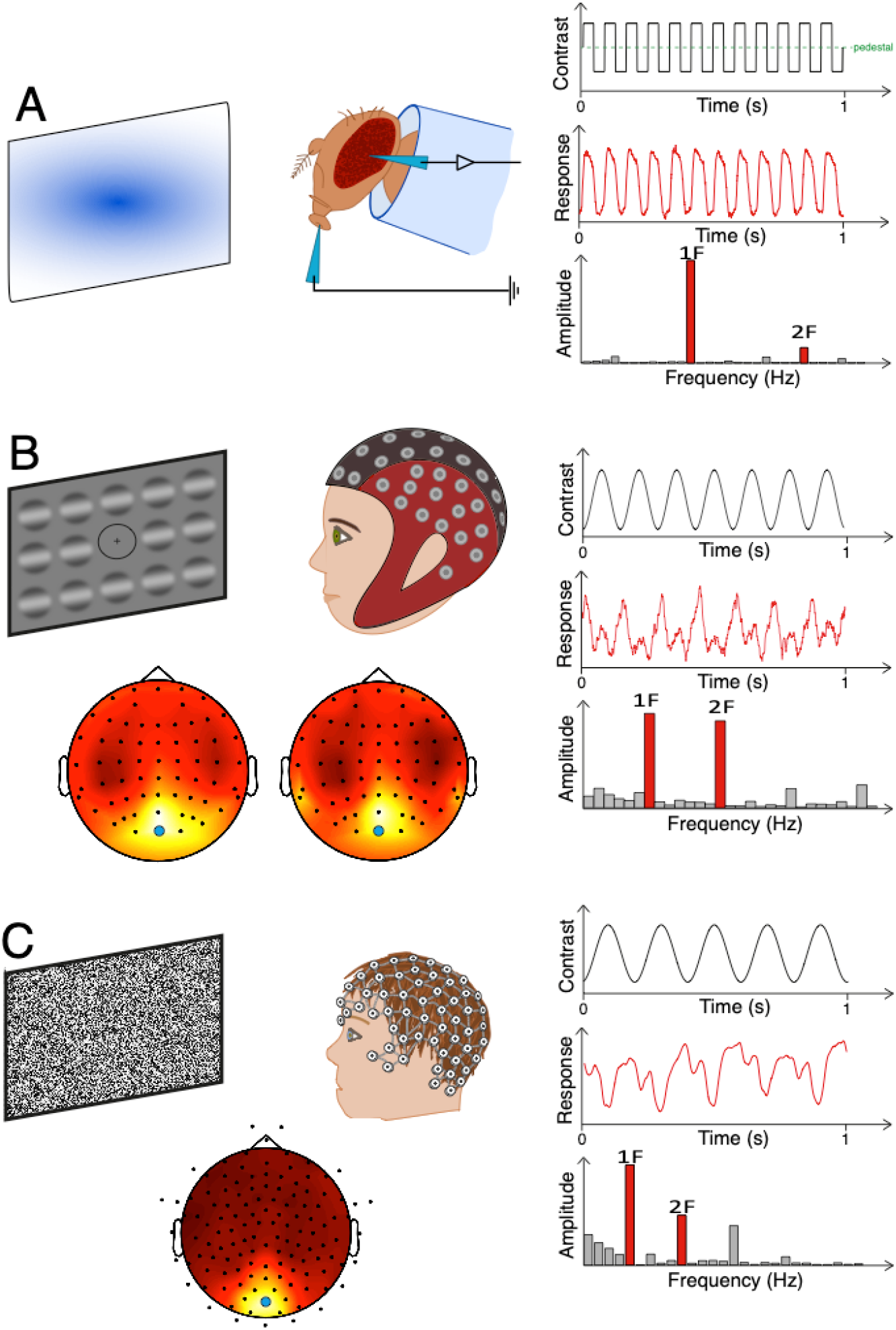
Human and *Drosophila* steady-state electrophysiology methods. Panel A left illustrates the experimental set up for fruit fly electrophysiology (see *Drosophila electroretinography* for more details). Panel A right shows the square wave stimulus trace flickering at 12Hz (top), example electrophysiological responses over time (middle) and Fourier-transformed response amplitudes in the frequency domain (bottom). Panel B left illustrates the experimental set up for adult participants, who were presented with a grid of sinusoidal gratings flickering at 7Hz whilst ssVEPs were recorded with a 64-channel EEG cap (top). SSVEPs were measured from occipital electrode Oz (blue circle) where the highest 1^st^ harmonic amplitude was centred (AQ adults – bottom left, ASD adults – bottom right). Panel B right shows the stimulus trace (top), example responses in the time domain (middle) and in the frequency domain (bottom). Panel C shows equivalent experimental set up, stimulus and response traces for the children’s dataset.

**Fig 2.**
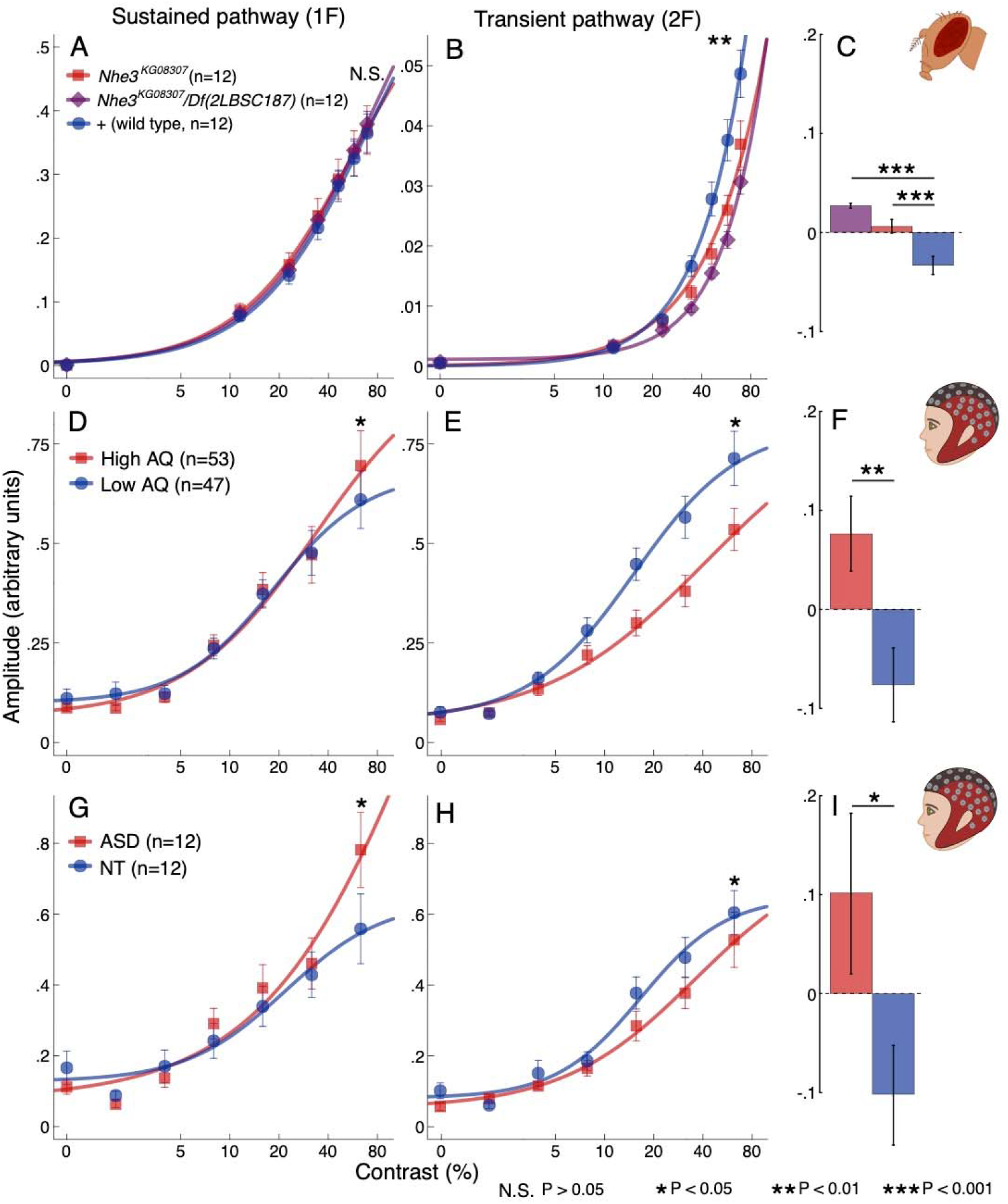
Older ASD-mimic flies and autistic humans show reduced visual responses in the transient component. Contrast response functions for adult *Nhe3* mutant flies (*Nhe3^KG08307^* homozygotes, red squares and *Nhe3^KG08307^ /Df(2L)BSC187*, purple diamonds) were similar at the first harmonic (a one-way ANOVA showed no effect of group F_2,33_ = 0.05, P = 0.95, panel A) but responses were reduced for P/P (simple contrast, P=0.025) and P/Df mutants compared to controls at the second harmonic (simple contrast P = 0.001; ANOVA group effect F_2,33_ = 6.71, P < 0.01; panel B). Ratios between frequencies 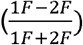 were significantly higher for P/P (P < 0.001) and for P/Df (P < 0.0001) than for the control genotype (C). First harmonic responses were also similar for the high AQ and low AQ groups (panel D) and for autistic and neurotypical adults (panel G). However, second harmonic responses were reduced for both adults with high AQ (panel E) and autistic adults compared to controls (panels H). The ratio between harmonics was also higher in both experimental groups compared to controls (panels F and I, P = 0.005 and P = 0.04, respectively). Curved lines are hyperbolic function fits to the data. Frequency ratios are baselined in respect to the mean over groups of each comparison for display purposes. Error bars in all panels represent ±SEM.

To quantify this functional dissociation whilst controlling for overall responsiveness of the visual system, we calculated a normalised ratio between first (1F) and second (2F) harmonics 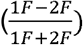 and averaged over the highest contrast conditions (where the response rises above the noise floor, see *Methods*). This allowed us to measure the differences between sustained and transient responses whilst normalising for overall responsiveness of the visual pathway. The ratio was significantly higher in both mutant strains than in the controls (ANOVA, F_2,33_ = 20.53, P < 0.0001, both paired contrasts P < 0.001; *Fig 2c*). These data suggest an impairment in the post-receptoral neural structures (downstream of the photoreceptors) of the older mutant flies (36).

Interestingly, unlike the older flies, the young 3-day-old flies showed a reduced response at both frequencies (see *Fig 3a,b*) relative to controls. Importantly, there was no effect of genotype on the ratio between harmonics (F_2,33_ = 1.38, P = 0.27; *Fig 3c*). These results suggest a deficit in the sustained visual module of young mutant flies. These differences between visual responses at two stages of life suggest a change in visual processing over the course of development.

**Fig 3.**
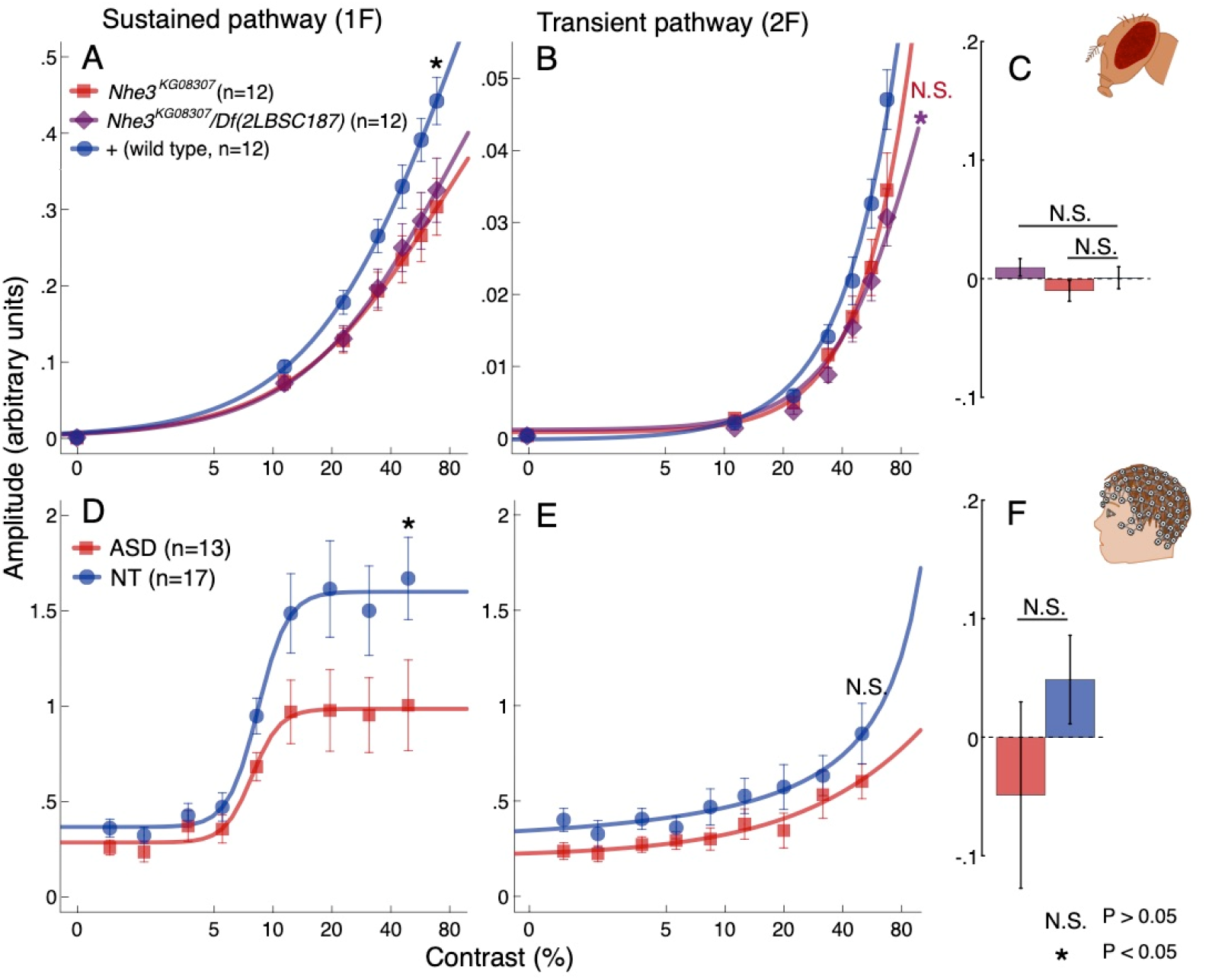
Young ASD-mimic flies and autistic children show reduced visual responses in the sustained component. Young fruit flies showed reduced responses at the first harmonic (F_2,33_ = 3.73, P = 0.035; panel A) with P/P and P/Df flies showing a significant difference from control flies (respectively, P = 0.016 and P = 0.040). There was also a significant effect of genotype at the second harmonic (F_2,33_ = 3.39, P = 0.046, panel B). P/Df flies showed a significant difference from control flies (P = 0.018), however, P/P showed a non-significant difference from controls (P = 0.064). The flies had normal frequency ratios (panel C). Autistic children also showed reduced first harmonic (t_28_ = 2.065, P = 0.048; panel D) but not second harmonic responses (t_28_ = 1.26, P = 0.22; panel E) and had frequency ratios similar to that of control children (t_28_ = 1.21, P = 0.24; panel F). Curved lines are hyperbolic function fits to the data. Frequency ratios are baselined in respect to the mean over groups of each comparison for display purposes. Error bars in all panels represent ±SEM.

### High autistic trait population show similar ssVEPs to *Nhe3* flies

To assess the relevance of the *Nhe3* model to the human ASD phenotype we used a comparable and similarly sensitive ssVEP paradigm in human participants. One hundred neurotypical participants with putative autistic traits measured using the Autism Spectrum Quotient (AQ) questionnaire (37) were tested with the ssVEP paradigm. Visual responses were recorded from an occipital electrode (Oz, located at the back of the head over the visual cortex) to grating stimuli flickered at 7Hz. Seven contrast conditions (each repeated eight times) were presented in a randomised order. First and second harmonic ssVEP responses were again derived via Fourier analysis. The evoked response data were averaged separately over participants split by their median (median = 14) AQ score: high (n = 53, AQ mean = 20.57, SD = 6.66) and low (n = 47, AQ mean = 9.47, SD = 3.08) AQ (high AQ implying many autistic traits). The second harmonic was notably reduced in the high AQ group, similarly to mutant fruit flies (*Fig 2d, 2e*). In addition, the first harmonic response was slightly increased in the high AQ group. A two-way ANOVA showed the interaction between group and frequency to be significant (F_1,98_ = 6.17, P = 0.015). The high AQ group also had a significantly higher frequency ratio than the low AQ group (t_98_ = 2.86, P < 0.01, *Fig 2f*). Moreover, a regression analysis showed that AQ scores correlated with the frequency ratio, with high AQ scores being predictive of higher ratios (R = 0.26 F_1,98_ = 6.87, P = 0.01; see *Fig 4*). This result shows a relationship between the amplitude of the second harmonic response and the severity of the subclinical ASD phenotype, however, this effect cannot be directly generalised to clinical autism as the AQ is not diagnostic of full-blown ASD.

**Fig 4.**
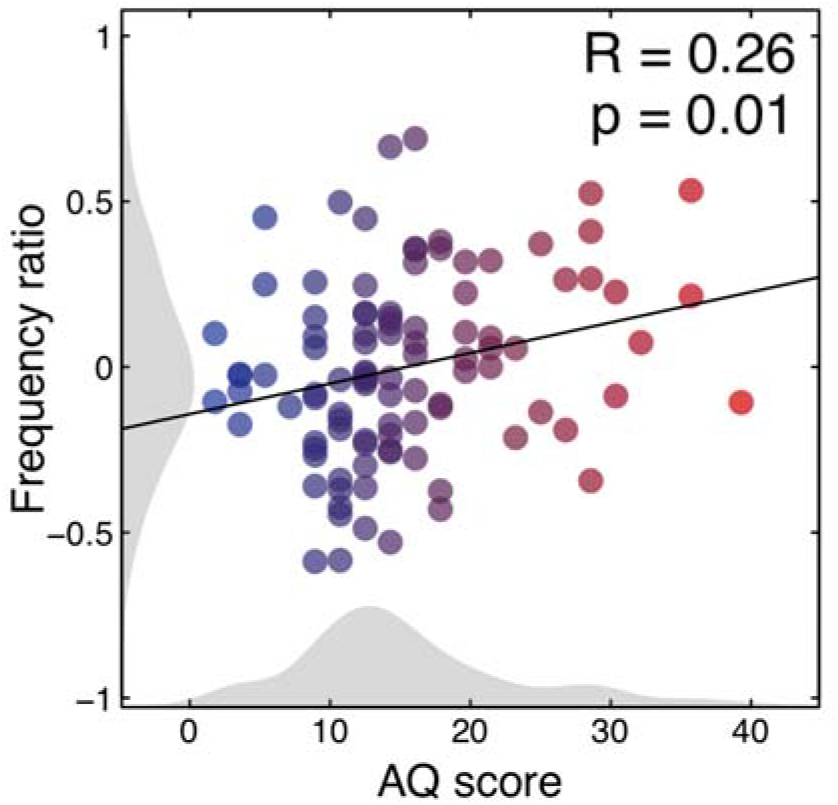
Positive relationship between the number of autistic traits and first/second harmonic ratio. Scatterplot showing a significant positive relationship between AQ scores and frequency ratios in the 100 neurotypical adult dataset indicating a gradual increase in response differences with the number of reported autistic traits. The black line indicates the regression line of best fit. Shaded grey areas show histograms of AQ scores and frequency ratios. Blue-red colour transition indicates number of AQ traits with participants split by median into low and high AQ groups as presented in *Fig 2*.

### Adult autistic individuals show a similar pattern of responses as mature *Nhe3* flies

We assessed the ssVEP difference between harmonics in clinical ASD by testing 12 typical-IQ autistic adults (diagnosis confirmed with the Autism Diagnostic Observation Schedule, Second Edition (ADOS-2), Lord et al., 2000) and 12 age- and gender-matched controls using the same human ssVEP paradigm. The pattern of data again mimicked that of the previous adult dataset: there was a significant interaction between group and frequency (F_1,22_ = 5.85, P = 0.02; *Fig 2g, 2h*), with the difference in second harmonic responses replicating that of the high AQ individuals and older mutant fruit flies. The ratio between harmonics was again significantly larger in the ASD group than in the control group (t_22_ = 2.13, P = 0.04; *Fig 2i*).

### Young *Nhe3* fly responses are similar to autistic children’s responses

Considering the striking similarity between the adult human datasets and the adult fruit fly model, it is reasonable to ask if similarities also exist between human children and young ASD-mimic flies. Specifically, our fly model predicts that the visual system of autistic children should show reduced responses at both the first and second harmonics. To examine this, we recorded from 13 autistic children (5 – 13 years old) and 17 neurotypical age-and gender-ratio-matched controls using an ssVEP contrast-sweep paradigm. Artifact rejection was employed to control for movement and blinking in both groups. The stimulus in each sweep trial increased continuously in contrast from 0% to 50% in logarithmic steps. Data were binned into 9 contrast levels before being Fourier transformed to compute response amplitudes.

As predicted by the model, the ASD group showed reduced amplitudes of the 1F, sustained response (t_28_ = 2.07, P = 0.04; *Fig 3d, 3e*) which was not found in the autistic adults, individuals with high AQ or older mutant fruit flies. A two-way ANOVA also revealed a significant group effect over both frequencies (F_1,28_ = 4.23, P = 0.049). Unlike adults, children exhibited no difference in frequency ratios between the groups (t = 1.41, P = 0.17; *Fig 3f*). Although children showed reduced amplitudes in the sustained response as predicted by the *Drosophila* model, the amplitude reduction observed in the fruit fly second harmonic responses was in the same direction, but was not statistically reliable in the children (t_28_ = 1.26, P = 0.219). This may be due to difficulty in measuring the relatively smaller F2 response in children.

## Discussion

We found sensory processing alterations in our *Drosophila* model of ASD that were consistent with similar response alterations in human data at two stages of development. Our steady-state electrophysiology data showed a selective depression in second harmonic visual responses in autistic adults, individuals with high levels of autistic traits and *Nhe3* mutant fruit flies, suggesting that this response alteration is specific to the autistic phenotype in mature individuals of both species. These differences were also present when we calculated 1^st^/2^nd^ harmonic ratios in order to control for changes in overall visual sensitivity. This suggests that the transient component of visual processing is selectively affected. Autistic children and young *Nhe3* flies showed an alteration in sustained visual processing, not present in the adults. The *Nhe3* fruit fly model of autism was predictive of these sustained visual response alterations both in children and in adults (atypical in early life, normal in later life), suggesting a fundamental and pervasive change in visual processing occurs during development in ASD. Although the human *Nhe9* is only one gene implicated in ASD, its ortholog in fruit flies was able to produce a measureable sensory processing effect, which has a close counterpart in human ASD.

We replicated the response alterations of autistic adults in neurotypical individuals with high AQ: this group had visual responses consistent with those of autistic participants diagnosed ASD suggesting common visual response properties between samples. This was unsurprising as previous research has found that AQ scores in the general population are highly correlated (R = 0.77) with sensory processing difficulties, as measured by the Glasgow Sensory Questionnaire (39), indicating that high AQ individuals exhibit milder forms of sensory difficulties.

The intact first harmonic response in adult flies and humans indicates normal functioning of mechanisms which give rise to the sustained response. Conversely, the reduced second harmonic response as well as the increased ratio between harmonics suggest a modification in the transient dynamics of the visual system. In fly, the first harmonic has been associated with sustained photoreceptor polarisation and the second harmonic with second-order lamina cells (31). In human, an association has been made between simple cell and sustained responses to pattern onset and between complex cells and transient responses at both stimulus onset and offset (40). Although simple cells exhibit some transient response properties as well (40,41), the intact first harmonic of adults suggests that their response modification is specific to human complex cells that only generate even-order response components. This early, cell-type-specific deficit may explain previous findings of atypical neural dynamics of spatial frequency processing in ASD in the face of normal sensitivity thresholds (20,42).

Mechanistically, lower 2^nd^ harmonic responses could either be generated by disturbances in non-linear transduction of visual signals or by subsequent temporal processing. As the 2^nd^ harmonic, by definition, has a higher temporal frequency, a bandpass temporal filter shifted towards lower frequencies would attenuate signals at this frequency more compared to the 1^st^ harmonic. There is at present no consistent evidence for lowered temporal resolution/prolonged integration in human ASD. One study found no difference between autistic and neurotypical participants (Kwakye et al 2011), one study found finer/higher temporal resolution (Falter et al., 2012) and another coarser/lower temporal resolution (de Boer-Schellekens, 2013). The possible role of temporal integration time could be tested in future work by using a lower stimulus frequency (such that the 2^nd^ harmonic would now equal the current 1^st^ harmonic frequency) and by observing whether the difference between harmonics disappears. An absence of a difference would indicate that temporal filtering is affected in ASD, whereas a persistently reduced 2^nd^ harmonic would indicate a difference in the non-linearity.

The differences in sustained and transient modules observed in our *Nhe3* model mimics the alteration of neural dynamics in autistic adults. *Nhe3* affects the exchange of sodium and hydrogen ions in cell membranes directly affecting neural signalling (33,43). Differential expression of *Nhe3* and other genes in ASD, which has been observed in other parts of the brain (33,44) may extend to differential expression in colour and motion modules in the *Drosophila* visual system. As *Nhe3* (*SLC9A9* in humans) is only a single gene in a multifaceted genetic etiology of autism, it is likely that the expression of several genes in human autism affects simple and complex cell dynamics, producing similar effects at the neural population level. Furthermore, such abnormality in gene expression in other parts of autistic brains, as well as environmental influences and gene-environment interactions, may give rise to a wide range of cognitive and social differences in childhood and adulthood.

Our data indicate little or no over-responsivity in the visual responses that are predicted by excitation/inhibition (E/I) imbalance theories (45,46) and consistent with measurements of some previous studies (47,48). However, it is possible that an E/I imbalance in autism stemming from GABA-ergic mechanism differences affects different neuron types or processing pathways in distinct ways and to different extents. It is also possible that E/I imbalance in sensory cortical areas in autistic individuals compensates for lower sensory signals (such as the second harmonic response here) in childhood. Regardless, cell-type based processing modifications may explain previous inconsistencies in studies of sensory symptoms in ASD that did not differentiate the relevant neural dynamics (17). Furthermore, the current results can provide an amended explanation to the magnocellular (M pathway) dysfunction hypothesis (16). As it is difficult to isolate the M pathway by changing stimulus properties (34), the paradigms previously used to investigate magnocellular dysfunction in ASD may have been selectively activating responses of transient components rather than the M pathway, in particular (16,49).

Developmentally, the observed lessening of the response modifications in both species with increasing age is in accordance with previous findings showing reduction or complete rescue of neuroanatomical differences present in early ASD childhood over the course of maturation (50). Previous longitudinal research has also shown that symptom severity in individuals diagnosed with ASD in childhood decreases over time (51,52). McGovern & Sigman (52) found that 48 adolescents, who were diagnosed with ASD as children, showed marked improvement in social interaction, repetitive/stereotyped behaviours and other symptoms, with two no longer meeting criteria for ASD under ADI-R criteria, and four under ADOS criteria. This might be explained by a change in neural processing during development, which would likely affect both complex behavioural and simpler sensory outcomes.

One possible mechanism that would explain the developmental change is that the atypical nature of neural signalling (such as ion balance in the case of *Nhe3*), changes over time. In flies, reduced *Nhe3* expression may reduce the rate at which sodium ions and protons are exchanged across the cell membrane. At least in mosquito, this exchanger is found in the gut, and Malpighian tubules (the fly equivalent of the kidney) (53). Failure to properly regulate ionic balance in young adult flies might affect the sodium concentration, or proton levels in the body and brain, and affect the speed and intensity of action potentials. Later in life, the normal balance may be restored. A similar reduction in efficacy of *SLC9A9*, linked to ASD, may also be present and explain the homology. In this respect, we note that another transporter, the potassium/chloride exchanger, has been linked to epilepsy in young people: with age the kcc/KCC2 eventually achieves a normal ionic balance and proper inhibitory GABA signalling (54).

The *Nhe3* model may facilitate further research on the development of ASD in young brains as well as the development of early biomarkers and treatments. Consistency between the fly and human datasets at both ages indicates a modification of a fundamental sensory mechanism comprising two components that have been conserved over 500 million years of evolution. The conservation of the phenotype and mechanisms from fly to human opens up the option to utilise the unrivaled genetic tractability of the fly to dissect the molecular mechanisms underpinning the disorder.

## Methods

### *Drosophila* stocks

Two *Drosophila melanogaster* genotypes were used as ASD models. The *Nhe3* loss-of-function P-element insertion (*Nhe3^KG08307^* homozygotes) mutation was homozygous *P{SUPor-P}Nhe3^KG08307^* (Bloomington Drosophila Stock Center (BDSC) 14715). The deficiency was Df(2L)BSC187 (BDSC 9672). To avoid second site mutations in the P-element stock, we used the hemizygote *Nhe3^KG08307^ /Df(2L)BSC187* as a second experimental genotype.

For our control cross we mated the lab stock of *Canton-S (CS*) flies with those with isogenic chromosomes 2C and 3J (55). All tested flies had dark red eyes. All genotypes were raised in glass bottles on yeast-cornmeal-agar-sucrose medium (10g agar, 39g cornmeal, 37g yeast, 93.75g sucrose per litre). They were kept at 25°C on a 12 hour light-dark cycle. Male flies were collected on CO_2_ the day after eclosion and placed on Carpenter (1950) (56) medium in the same environmental conditions for either 3 days or 14 days. Flies were tested approximately between the 4^th^ and 9^th^ hour of the daylight cycle.

### *Drosophila* electroretinography

Steady-state visual evoked potentials (SSVEPs) were obtained from the fruit flies (25,31). Flies were recorded in pairs in a dark room. They were placed in small pipette tips and secured in place with nail varnish. One glass saline-filled electrode was placed inside the proboscis of the fly and another on the surface of the eye. A blue (467nm wavelength) LED light (Prizmatix FC5-LED) with a Gaussian spectral profile (FWHM 34nm) was placed in front of the flies together with a diffuser screen and used for temporal contrast stimulation. Flies were dark adapted for at least two minutes and then tested for signal quality with six light flashes. Steady-state stimulation lasted 12 min and comprised seven contrast levels (0 – 69% in linear steps) each with five repetitions. The frequency of the light flicker was 12Hz. Each trial (contrast level repetition) was 11 s. The order of the contrast conditions was randomised. The stimulation and the recording from the fly was controlled by in-house MATLAB scripts (scripts can be found in https://github.com/wadelab/flyCode).

### Adult EEG

One-hundred neurotypical adult participants (32 males, mean age 21.87, range 18 – 49, no reported diagnosis of ASD, reportedly normal or corrected to normal vision) took part in the autism spectrum quotient (AQ) measurement study. The AQ is an instrument used for quantifying autistic traits in the neurotypical population and has been shown to have high face validity and reliability in these populations (37). Due to time constraints we used an abridged version of the AQ questionnaire which consists of 28 questions rather than the typical 50 (AQ-Short, (57)). Scores were then scaled to fit the conventional AQ scale. Each participant completed the AQ questionnaire on a computer in the laboratory. The participants were then median split (median = 14) into high and low AQ groups.

For the autistic adult ssVEP study, 12 typical-IQ autistic participants and 12 gender- and age-matched controls (11 males, mean age 23.53, range 18 – 39, reportedly normal or corrected to normal vision) took part. ASD diagnosis was confirmed with the Autism Diagnostic Observation Schedule, second edition (ADOS-2). Although IQ was not explicitly measured in this study, all adults had normal speech and a high level of independence (the majority were university students). The absence of ASD diagnosis in the neurotypical participants was also confirmed with ADOS-2 (none of the control participants met criteria for ASD). All participants in the study gave informed consent and were debriefed on the purpose of the study after the experiment. The experiments were approved by the Department of Psychology Ethics Committee at the University of York.

Steady-state VEPs were recorded using an ANT Neuro system with a 64-channel Waveguard cap. EEG data were acquired at 1kHz and were recorded using ASALab, with stimuli presented using MATLAB. The timing of the recording and the stimulation was synchronised using 8-bit low-latency digital triggers. All sessions were performed in a darkened room, testing lasted 45-60min with approximately 20min set up time.

Stimuli were presented on a ViewPixx display (VPixx Technologies Inc., Quebec, Canada) with a mean luminance of 51cd/m^2^ and a refresh rate of 120Hz. Stimuli were 0.5 cycle/deg sine-wave gratings enveloped by a raised cosine envelope. Gratings subtended 3 degrees of visual angle and were tiled in a 17×9 grid. The participants fixated on a circle in the middle of the screen and performed a fixation task (two-interval-forced-choice contrast discrimination) to maintain attention. All participants were able to perform the task at above chance levels. There were seven contrast conditions for the flickering gratings (0%, and 2 - 64% in logarithmic steps, where C_%_ = 100(*L_max_*−*L_min_*)/(*L_max_*+*L_min_*), *L* is luminance) and eight repetitions. Stimuli flickered on/off sinusoidally at 7Hz. Trials were presented in random order in four testing blocks with short breaks in between. Each trial was 11 seconds long and contained gratings of a random spatial orientation to avoid orientation adaptation effects. These trials were intermixed with orthogonal masking trials that are not presented as part of this study. Data were taken from the occipital electrode Oz.

### Child EEG

Thirteen children with a diagnosis of ASD and 20 neurotypical controls matched on gender ratio (10 and 12 males respectively) and average age (mean age 9.31 and 8.94 respectively, range 5 – 13) completed the study. Three of the neurotypical children were tested but excluded due to having autistic siblings (17 participants were included). All children were in mainstream local schools (if they were old enough) and did not have other (or any – in the case of the neurotypical group) reported history of serious medical, psychiatric, or neurological conditions.

Steady-state EEG data were acquired with a 128-channel HydroCell Geodesic Sensor Net (Electrical Geodesics Inc.). Data were digitised at 432Hz and band-pass filtered from 0.3Hz to 50Hz and were recorded using NetStation 4.3 Software. Highly noisy data were excluded by removing repetitions with amplitudes that were four standard deviations away from the group mean (for each contrast level and harmonic individually). There were 10 repetitions in total, however, two autistic and one neurotypical child only completed 8 repetitions.

Increasing contrast sweep ssVEPs were used. Stimuli for this experiment were presented on an HP1320 CRT monitor with 800×600 pixel resolution, 72Hz refresh rate and mean luminance of 50cd/m^2^. Stimuli were random binary noise patterns of two luminance levels that increased in contrast in 9 logarithmic steps (0% – 50%) of 1 second each. Each trial contained a prelude at the initial value of the sweep and a postlude at the final sweep value, lasting 12 seconds in total. Stimuli flickered at 5.12Hz. Data from the middle 9 seconds during the sweep were binned according to contrast steps. Methodological differences between the adult and child datasets were due to different conventions being used by the two laboratories in which data were collected.

### Data analysis

A Fast Fourier transform (in MATLAB) was used to retrieve steady-state response amplitudes at the stimulation frequency (12Hz for fruit flies, 7Hz for adult participants and 5.12Hz for children) and at the second harmonic (24Hz, 14Hz and 10.24Hz respectively). Fourier transforms were applied to 10 s of each trial (first 1s discarded; total trial length was 11s) for the fruit fly and the adult participant datasets and to 1 second binned data for the children’s dataset. Contrast response functions were obtained by coherently averaging the amplitudes over repetitions for each contrast level within a participant. Group/genotype scalar means over response amplitude (discarding phase angle) were then calculated for each contrast across participants/flies.

Two-way (harmonic x group) ANOVAs were performed on amplitudes at the highest contrast level to investigate the interactions and group effects in all human datasets where only two groups were compared. To identify at which harmonic the autistic children showed a decreased response, two independent samples t-tests were also conducted. One-way ANOVAs with simple planned contrasts were conducted to assess the genotype differences in fruit fly first and second harmonic responses separately as that aided the interpretability of the results between the three genotypes.

To investigate the dissociation between first and second harmonic responses a scaled ratio 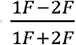 (where 1F is the first and 2F is the second harmonic) was calculated for each participant/fly and each contrast condition. To increase the power of statistical analyses and to decrease the type I error rate, the ratios were then averaged over the contrast conditions that had first harmonic amplitudes significantly above the baseline response (0% contrast condition). For fruit flies this was six conditions (11.5 – 69%), for adult participants this was four conditions (8 – 64%) and for children this was five conditions (8.5 – 50%). This procedure resulted in a single frequency-ratio index for each participant/fly. One-way ANOVAs with simple planned contrasts (comparing mutant genotypes with the control genotype) were conducted on the fly frequency ratios for each age separately. Independent t-tests were used to compare frequency ratios in all human datasets between groups. Additionally, a linear regression was conducted on the adult AQ measurement dataset to assess the predictive power of AQ scores on the ratios between frequencies. All statistical tests were two-tailed.

## Authors’ contributions

All authors contributed to conceiving and designing of experiments; G.V., F.P. collected data; G.V., D.H.B., A.M.N. performed statistical analyses; G.V., A.R.W., A.M.N., D.H.B. interpreted the results with contributions from all authors; G.V. wrote the manuscript with A.M.N, D.H.B. and A.R.W. with input from all authors.

## Acknowledgements

We would like to thank E. Cole for second-coding ADOS2 assessments; N. Gutmanis, S. Harris, R.E. Kitching, H.A. Melton, R. Norman, C.J.T. Scott, C. Simpson, A.K. Smith, A. Stockton and K. Wailes-Newson for contributing to EEG data collection. A.M.N. and F.P. were supported by a grant from the Simons Foundation Autism Research Initiative. We would also like to thank the Experimental Psychology Society for awarding G.V. the Grant for Study Visits for travel to Stanford University. Lastly, we would like to thank all our participants and their families for their contribution.

